# Protective role of N-acetylcysteine and Sulodexide on endothelial cells after SARS-CoV-2 infection

**DOI:** 10.1101/2023.07.19.549800

**Authors:** Justyna Rajewska-Tabor, Patrycja Sosińska-Zawierucha, Malgorzata Pyda, Andrzej Bręborowicz

**Author notes:** Corresponding author: Justyna Rajewska-Tabor, I Clinic of Cardiology, Department of Magnetic Resonance, Dluga 1/2, 61-848 Poznan, Poland.

## Abstract

Severe acute respiratory syndrome coronavirus-2 causes hyperinflammation and activation of coagulation cascade and in the result aggravates endothelial cell dysfunction. N-acetylcysteine and Sulodexide have been found to mitigate endothelial damage. The influence on coronary artery endothelial cells of serum collected after 4+/-1 months from coronavirus infection was studied. The concentrations of serum samples of interleukin 6, von Willebrand Factor, tissue Plasminogen Activator and Plasminogen Activator Inhibitor-1 were studied. The cultures with serum of patients after coronavirus infection were incubated with N-acetylocysteine and Sulodexide to estimate their potential protective role. The blood inflammatory parameters were increased in the group of cultures incubated with serum from patients after coronavirus infection. Supplementation of the serum from patients after coronavirus infection with N-acetylcysteine or Sulodexide reduced the synthesis of interleukin 6, von Willebrand Factor. No changes in the synthesis of tissue Plasminogen Activator were observed. N-acetylcysteine reduced the synthesis of Plasminogen Activator Inhibitor-1. N-acetylcysteine and Sulodexide increased the tPA/PAI-1 ratio. N-acetylcysteine may have a role in reducing the myocardial injury occurring in the post-COVID-19 syndrome. Sulodexide can also play a protective role in post-COVID-19 patients.

## INTRODUCTION

Severe acute respiratory syndrome coronavirus-2 (SARS-CoV-2) is a pandemic disease affecting the respiratory system and also other organs of the human body [1]. The virus invades the cells of the respiratory system, entering it mainly through endothelial cells, and then attacks other organs and endothelial cells themselves [2]. The action of the virus and the human defence mechanisms activate anti-viral processes, causing hyperinflammation, which then activates neutrophils, monocytes and platelets and results in the activation of the coagulation cascade, possibly leading to intravascular thrombosis [3,4,5].

At the cellular level, the structural and functional dysfunction of endothelium is due to the lack of nitric oxide (NO), cellular oxidative stress, the inflammatory process and a damaged glycocalyx structure. Myocarditis is one of the complications of COVID-19, which may aggravate endothelial cell dysfunction [2]. All the above and the resulting myocardial damage, acting on the endothelial cells of the coronary artery [6,7,8,] may be responsible for the long-COVID −19 syndrome, which affects 10-30% of patients [9,10].

N-acetylcysteine (NAC) and Sulodexide have been found to mitigate endothelial damage and dysfunction [11,12,13]. By suppressing the secretion of pro-inflammatory cytokines such as NF-κB, IL-8 and IL-6, NAC reduces the chemotactic migration of monocytes [14,15]. NAC has also been shown to have a protective role in reducing the replication of various viruses, including the human immunodeficiency virus (HIV) [16] or the respiratory syncytial virus (RSV) [17]. This protective effect of NAC on endothelial cells reduces the adverse effects of viruses on the vascular endothelium [18]. The results of earlier studies of other RNA viruses have suggested that NAC might also have a similar role in SARS-CoV-2 infection. NAC has also been shown to have the potential ability to inhibit SARS-CoV-2 replication [19,20,21,22]. Also, the supportive effect of Sulodexide in the acute phase of COVID-19 has been demonstrated in the literature [23,24,25]. The long-term protective effect of these substances is still being investigated [26].

We present results from the study in which effect of serum isolated from the postCOVID-19 patients on function of the human coronary endothelial cells was studied in in vitro culture. Additionally we evaluated if NAC and Sulodexide modify the post-COVID-19 serum-induced changes in these cells.

## MATERIAL AND METHODS

The effect of serum on the coronary endothelial cells collected from 12 patients with prior SARS-CoV-2 infection but otherwise healthy was evaluated in an in vitro culture. No patient showed a sign of the acute or persisted myocardial injury, as reflected by troponin I and NT-proBNP serum concentration. All patients had undergone a mild form of COVID-19, manifesting itself mainly in fever (9;75%), headache (8;67%), fatigue (7;58%) and muscle pain (7;68%). None of them was hospitalized during infection and had no need of oxygen treatment. They were isolated for at least 10 days with symptomatic treatment only. The laboratory tests after 4±1 months did not show any abnormalities: all patients had normal levels of D-dimers, C-reactive protein, creatinine and leukocytes.

The serum of 12 healthy individuals with no past COVID-19 experience was used as the control. The protocol of the study was approved by the Bioethical Committee of the Poznan University of Medical Sciences. All patients gave their written consent to participate in the study.

The biochemical characteristics of the studied groups are shown in Table 1. The concentrations of serum samples of interleukin 6 (IL6), the von Willebrand Factor (vWF), the tissue Plasminogen Activator (tPA) and Plasminogen Activator Inhibitor-1 (PAI-1) were studied with the commercially available ELISA kits (R&D Systems, Minneapolis, MN, USA).

**Table 1.** Characteristic of studied group and control group.

|  | Control | Post-COVID-19 group | p |
| --- | --- | --- | --- |
| Number of patients | 12 | 12 | - |
| Age (years) | 42.8±7.3 | 41.9±10.7 | ns |
| Time after COVID-19 (months) | 0 | 4.2±1.1 | <0.001 |
| Blood IL6 (pg/mL) | 3.5±1.3 | 45.6±90.9 | <0.001 |
| Blood vWF (µg/ml) | 5.6±0.9 | 41.1±6.6 | <0.001 |
| t-PA (mg/mL) | 3.7±1.5 | 34.8±16.6 | <0.001 |
| PAI-1 (ng/mL) | 18.9±2.6 | 74.1±13.3 | <0.001 |

IL6 – interleukin 6, vWF – von Willebrand Factor, t-PA – tissue Plasminogen Activator, PAI-1 – Plasminogen Activator Inhibitor-1. Correlation revised with Spearman tests.

### In vitro culture of the endothelial cells

During the experiments, the primary cultures of human coronary artery endothelial cells (CAEC) obtained from Cell Applications, Inc. (San Diego, California, USA) were used. The culture for the growth of the cells was provided by the producer. The cells were grown to monolayers in 75cm2 culture flasks, and were subsequently harvested with trypsin 0.05%EDTA 0.02% solution and seeded into the 48 wells culture plates. Experiments were performed on the endothelial monolayers.

### Effect the serum samples on the endothelial cells

The endothelial monolayers in 48 wells culture plates were exposed for 24 hours to the standard culture medium supplemented with the serum sample (20%v/v). In our earlier experiments we found that such treatment did not induce any morphological changes in the endothelial cells and while the MTT test (Abcam, Cambridge, UK) was used, it did not reduce viability either. The cells were exposed to MTT salt [3-(4,5-dimethylthiazol-2-yl)-2,5-diphenyltetrazolium bromide] for 3 hours at 37 °C. The generated formazan product was lysed and the lysate absorbance was measured at 595 nm.

The following experimental groups were studied, with cells exposed to the following solutions:

1. Culture medium (Control),
2. Culture medium supplemented with serum (20% v/v) from healthy donors,
3. Culture medium supplemented with serum (20% v/v) from the post-COVID-19 patients,
4. Culture medium supplemented with serum (20% v/v) from the post-COVID-19 patients + N-Acetylcysteine 1 mmol/L,
5. Culture medium supplemented with serum (20% v/v) from the post-COVID-19 patients + Sulodexide (0.5 LRU/mL).

### The cell parameters studied

#### Intracellular oxidative stress

At the end of the 24 hours incubation, in 6 wells from each group, the oxidative stress was measured. Free radicals generated within the cells were measured during their 45 minutes incubation at 37oC with a 2’7’dochlorodihydrofluorescein diacetate probe. After the lysis of the cells, the fluorescence of the cells lysates was measured in a fluorimeter at a wavelength of 485 nm for excitation and 535 nm for emission. The number of free radicals generated was expressed as a number of cells counted in 6 wells from each group, in separate wells.

#### Secretory activity of the cells

After 24 hours of incubation the medium in all wells was replaced with the standard culture medium for evaluation of the cell’s secretory activity in the following 24 hours. At the end of the incubation, the medium was collected from all wells, spun down (200g; 10 minutes) and frozen at −86°C for further analysis. The cells were harvested with a trypsin 0.05%EDTA 0.02% solution and counted in a hemocytometer. In the supernatants, the concentrations of the following molecules: IL6, tPA, PAI-1 and vWF were measured with the standard ELISA kits (R&D Systems, Minneapolis, MN, USA). The secretion of the molecules from the endothelial cells was expressed per number of cells.

#### Statistical analysis

The results are presented as a mean ± SD. The statistical analysis was performed with the Mann Whitney test, or ANOVA with the post hoc analysis of the Kruskal-Wallis test. The correlation between the studied groups was measured with the Spearman test. A p-value less than 0.05 was considered statistically significant.

## RESULTS

There was a significant difference between the blood inflammatory parameters which were increased in the post-COVID-19 group and remained unchanged in the control group (Table 1). The exposure of CAEC to the sera used in the experiments significantly modified the functional properties of the cells.

In the presence of the post-COVID-19 serum, the intracellular generation of free radicals was increased (+29%, p<0.001) in comparison with the cells exposed to the control medium (Figure 1). Supplementation of the post-COVID-19 serum with NAC reduced the intra-cellular oxidative stress (-23%, p<0.001). Post-COVID-19 serum stimulated the synthesis of IL6 in CAEC (+43%, p<0.001) as compared to cells treated with the control serum (Figure 1A). Supplementation of the post-COVID-19 serum with NAC or Sulodexide reduced the synthesis of IL6 to: −18%, p<0.02 and −24%, p<0.01, respectively (Figure 1A). In CAEC exposed to the post-COVID-19 serum, the synthesis of vWF increased by 22%, p<0.01, as compared to the control serum (Figure 1B). NAC used as a supplement to the post-COVID-19 serum reduced the synthesis of vWF by 30%, p<0.001 (Figure 2B). No changes in the synthesis of tPA in CAEC treated with the post-COVID-19 serum were observed (Figure 3A). However, the synthesis of PAI-1 was increased in the endothelial cells exposed to the postCOVID-19 serum (+20%, p<0.01) (Figure 3B). NAC reduced the stimulatory effect of the post-COVID-19 serum on the synthesis of PAI-1 in CAEC: 17%, p<0.002 (Figure 3B). The tPA/PAI-1 ratio, reflecting the net fibrinolytic activity of serum, was reduced in the postCOVID-19 group as compared to the control serum: 1.3±0.1 vs 1.6±0.1 (p<0.001). NAC and Sulodexide increased the tPA/PAI-1 ratio in the cells treated with the post-COVID-19 serum to 1.6±0.1 (p<0.01) and 1.5 ±0.1 (p<0.05), respectively.

**Figure 1.**
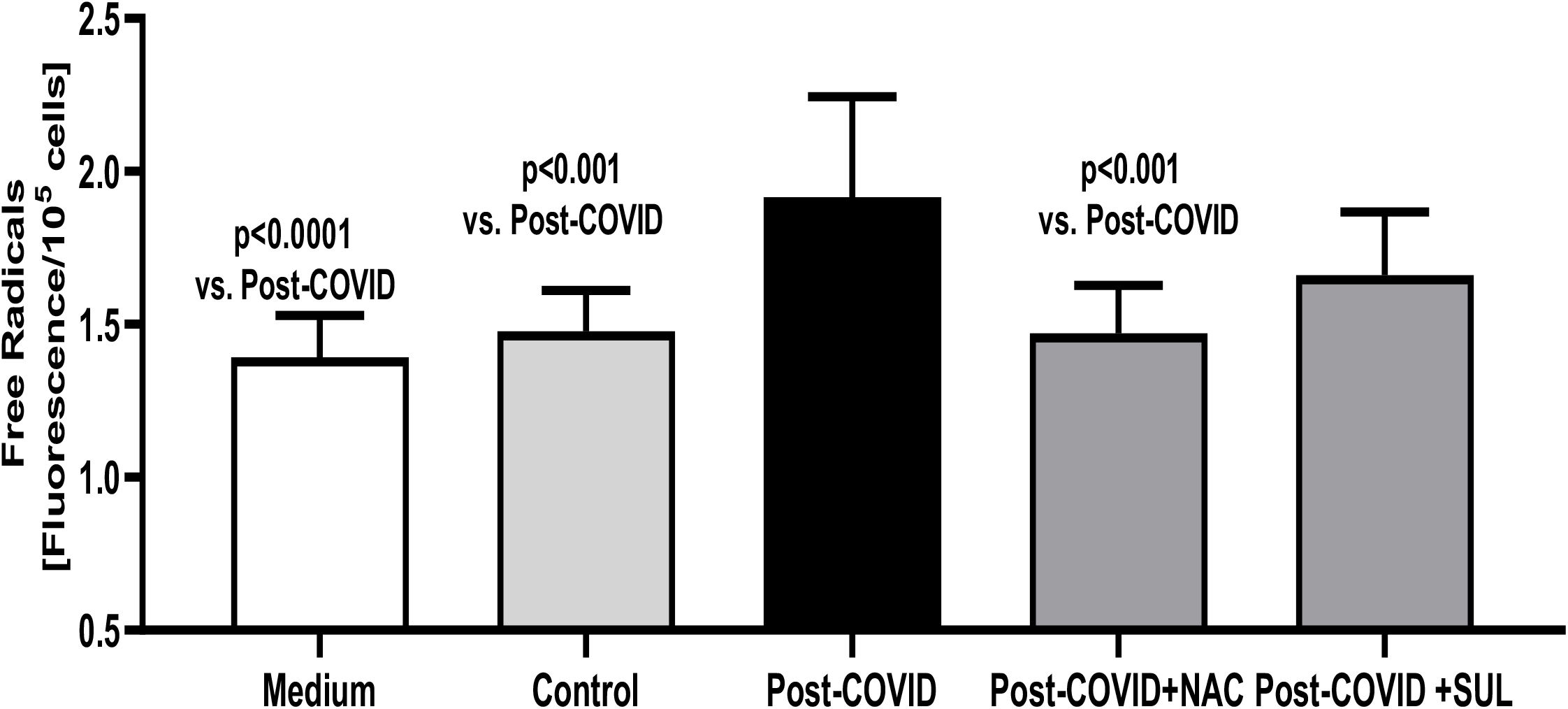
Intracellular generation of free radicals in CAEC exposed to culture medium (Medium), culture medium supplemented with 20% control serum (Control), 20% Post-COVID19-serum (Post-COVID), 20% Post-COVID-19 serum supplemented with N-Acetylcysteine 1 mmol/L (Post-COVID+NAC) or Post-COVID-19 serum with Sulodexide 0.5 LRU/mL (Post-COVID+SUL)

**Figure 2.**
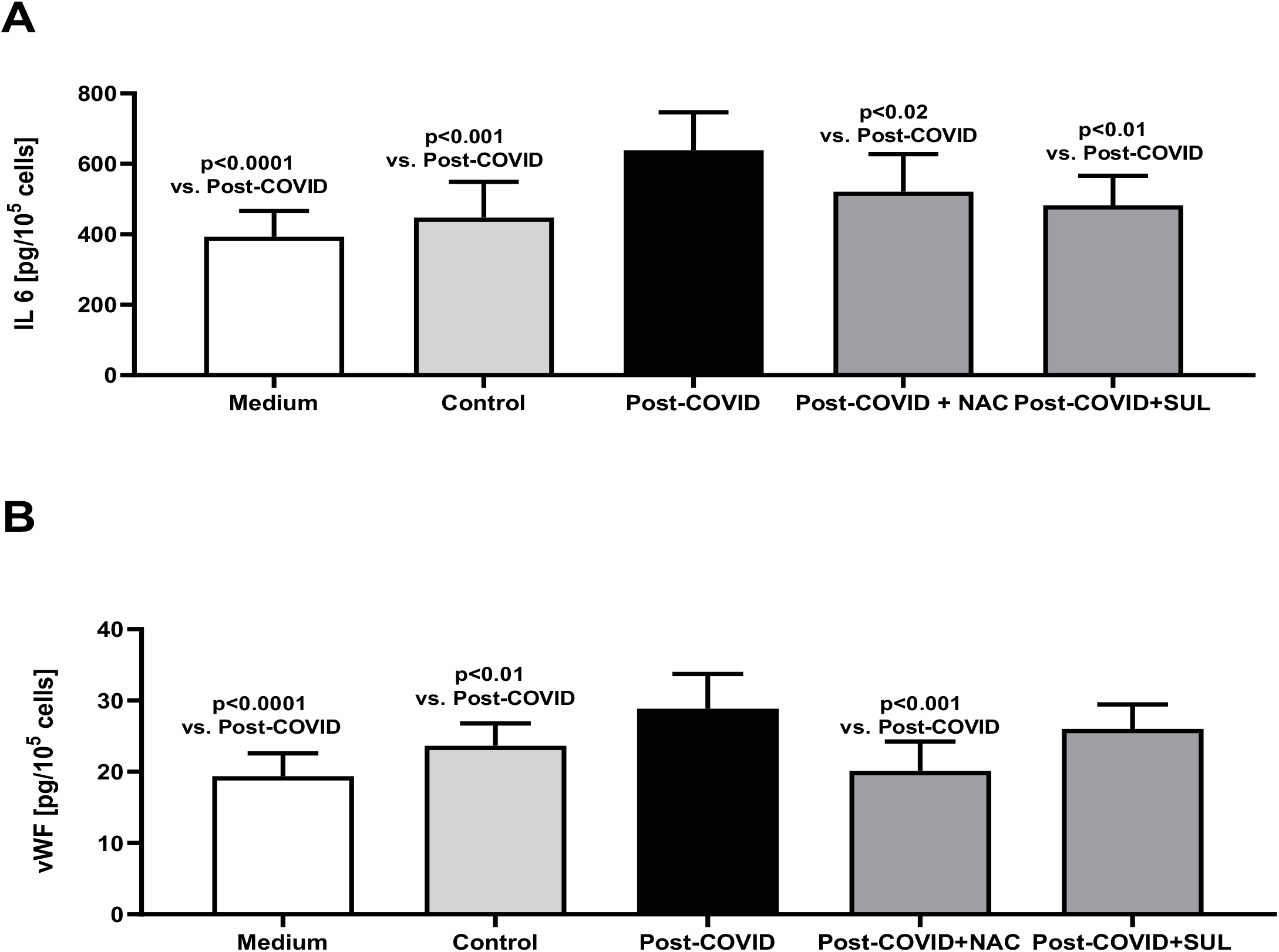
Synthesis of IL-6 (A), vWF (B), in CAEC exposed to culture medium (Medium), culture medium supplemented with 20% control serum (Control), 20% Post-COVID-19 serum (Post-COVID), 20% Post-COVID-19 serum supplemented with N-Acetylcysteine 1 mmol/L (Post-COVID+NAC) or Post-COVID-19 serum with Sulodexide 0.5 LRU/mL (Post-COVID +Sul)

**Figure 3.**
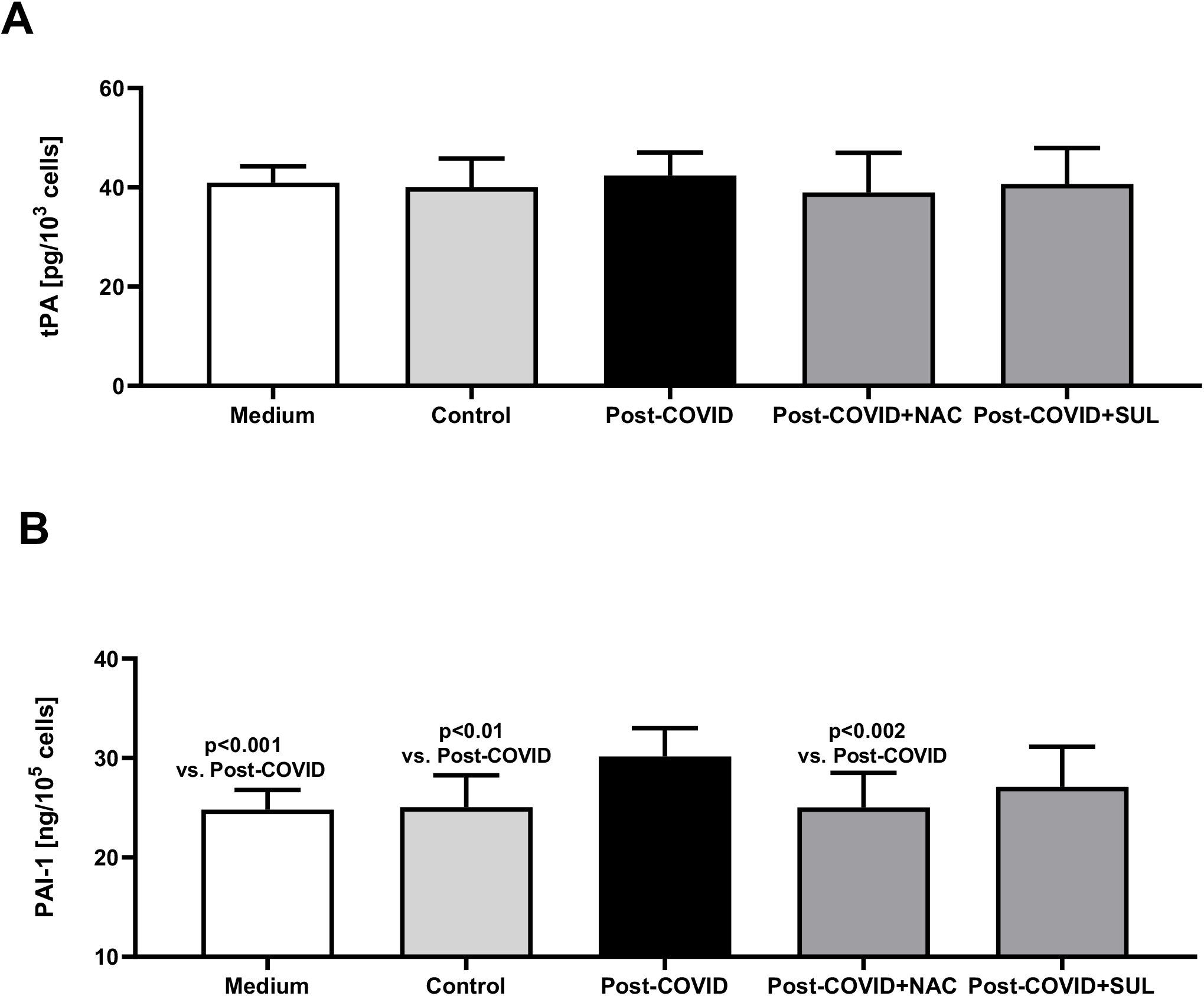
Synthesis of tPA (A), PAI-1 (B), in CAEC exposed to culture medium (Medium), culture medium supplemented with 20% control serum (Control), 20% Post-COVID-19 serum (Post-COVID), 20% Post-COVID-19 serum supplemented with N-Acetylcysteine 1 mmol/L (Post-COVID+NAC) or Post-COVID-19 serum with Sulodexide 0.5 LRU/mL (Post-COVID +Sul)

## DISCUSSION

The post-COVID-19 syndrome and its long-term effects, in particular, are still poorly understood and affect many areas and organs, showing a broad spectrum of symptoms. An injury of endothelial cells acquired during COVID-19 may have long-lasting consequences. Haffke M et al. have proved that the serum markers of an endothelial injury may be elevated as late as eight months after COVID-19 or even later [27]. In this study, we have proved that the inflammation of endothelial cells in SARS-CoV-2 infection may not only be caused directly by the virus but may also develop much later since the serum of the convalescents retains the possibility of inducing inflammation even months after infection. The exposure of CAEC to post-COVID-19 serum generates oxidant stress and stimulates CAEC to produce pro-inflammatory cytokines: IL6, vWF, tP, and PAI-1.

The thromboembolic complications of COVID-19 can be monitored by comparing the elevated levels of D-dimers, the fibrinogen and the vWF with the normal ranges of PT, aPTT and platelet count [28,29]. As reported in the literature, endothelial dysfunction biomarkers such as vWF and PAI-1 are increased in COVID-19 patients compared to healthy subjects. They seem to have a prognostic significance, being associated with more severe forms of the disease and a high mortality [30,31]. However, it has also been proved that sometimes thromboembolic complications develop in COVID-19 patients with normal PT, aPTT, aPTT and platelet results. In such an event, they may be monitored by checking the D-dimer, fibrinogen or vWF levels. Fogarty et al. showed sustained endotheliopathy measured by vWF level at a median of 68 days following SARS-CoV-2 infection [8]. This experiment revealed that there still occurs endothelial damage and increased secretion of endothelial factors such as vWF and PAI-1 four months after infection.

It has been reported that NAC decreases the binding of the virus to cells, decreases virus replication, has anti-inflammatory and antioxidant activity and modulates the immune system [21,32,33]. Therefore NAC is used in treating the acute period of SARS-CoV-2 infection, mainly intended to combat the cytokine storm [32,33].

Our experiment demonstrated the protective role of NAC for coronary artery endothelial cells. When exposed to post-COVID-19 serum and treated with NAC, CAEC released lower levels of pro-inflammatory cytokines than when exposed to post-COVID-19 serum without the NAC treatment. Moreover, we also proved that the incubation of endothelial cells with the serum of post-COVID-19 patients and supplemented with NAC resulted in the reduction of the secretion of the tested substances to the level of the CEAC secretion of the serum of healthy subjects.

Sulodexide is another possible candidate for application in COVID-19 therapy, particularly in patients with a mild form of the disease [23,26]. Sulodexide is a mixture of glycosaminoglycans consisting of 20% of dermatan sulfate and 80% fast-moving heparin. Its *in vitro* effects are comparable to enoxaparine, at least in anti-hemostatic effects [34]. Sulodexide produces multifaceted effects: by increasing tPA production and inhibiting platelet aggregation it activates arterial and venous anticoagulant and fibrinolytic processes, and it shows an anti-inflammatory activity, including the inhibition of IL-6 production [13,34]. It may be applied to treat different types of endothelial cells, as has already been demonstrated in other studies [11,13]. What is essential, Sulodexide may also be used in patients with renal impairment and is less likely associated with bleeding risk and heparin-induced thrombocytopenia.

In studies by Gonzalez-Ochoa AJ et al., when compared to a placebo, Sulodexide has been found that it may reduce the risk of hospitalization and the need for oxygen supply while improving laboratory parameters without increasing the risk of bleeding in early high-risk COVID-19 patients [25]. In this experiment, we have demonstrated that Sulodexide inhibits IL-6 secretion from endothelial cells at a later stage of COVID-19. The effects on other factors that we also tested are less significant. Despite this tendency to decrease the secretion of tPA, vWF and PAI, no statistically significant decrease in these factors has been determined. This finding, however, requires further investigation.

In conclusion a risk of a potential injury of endothelial cells remains months after COVID-19. NAC may have a role in reducing the myocardial injury occurring in the postCOVID-19 syndrome by reducing the endothelial injury of coronary arteries. Likewise, Sulodexide may also play a particular role in protecting endothelial cells in patients with or after COVID-19 infection

